# Low genetic variation in tolerance to defoliation in a long-lived tropical understorey palm

**DOI:** 10.1101/493932

**Authors:** Merel Jansen, Pieter A. Zuidema, Aad van Ast, Frans Bongers, Marcos Malosetti, Miguel Martínez-Ramos, Juan Núñez-Farfán, Niels P.R. Anten

## Abstract

Defoliation is a ubiquitous stressor that can strongly limit plant performance. Tolerance to defoliation is often associated with compensatory growth. Genetic variation in tolerance and compensatory growth responses, in turn, play an important role in the evolutionary adaptation of plants to changing disturbance regimes but this issue has been poorly investigated for long-lived woody species. We quantified genetic variation in plant growth and growth parameters, tolerance to defoliation and compensatory responses to defoliation for a population of the understorey palm *Chamaedorea elegans*. In addition, we evaluated genetic correlations between growth and tolerance to defoliation.

We performed a greenhouse experiment with 731 seedlings from 47 families with twelve or more individuals of *C. elegans*. Seeds were collected in southeast Mexico within a 0.7 ha natural forest area. A two-third defoliation treatment (repeated every two months) was applied to half of the individuals to simulate leaf loss. Compensatory responses in specific leaf area, biomass allocation to leaves and growth per unit leaf area were quantified.

We found that growth rate was highly heritable and that plants compensated strongly for leaf loss. However, genetic variation in tolerance, compensation, and the individual compensatory responses was low. We found strong correlations between family mean growth rates in control and defoliation treatments. We did not find indications for growth-tolerance trade-offs: genetic correlation between tolerance and growth rate were not significant.

The low genetic variation in tolerance and compensatory responses observed here suggests a low potential for evolutionary adaptation to changes in damage or herbivory, but high ability to adapt to changes in environment that require different growth rates. The strong correlations between family mean growth rates in control and defoliation treatments suggest that performance differences among families are also maintained under stress of disturbance.

## Introduction

Defoliation due to herbivory, pathogens, physical damage or harvesting is an ubiquitous stressor that can strongly limit individual plant performance (*i.e*. growth, reproduction and survival) as it entails a reduction in photosynthesis and resources, and thus in future growth. Performance reductions due to defoliation are often proportionately smaller than expected based on the fraction of leaf area that is being removed [1, 2] and in some cases plants even increase their performance under defoliation [3, 4]. In that sense plants can be tolerant to defoliation, and this tolerance is often associated with compensatory growth, a mechanism by which negative effects of leaf loss are mitigated [5]. There are three types of compensatory growth responses: plants can compensate for growth by allocating more new assimilates to leaves, by allocating new assimilates more efficiently to leaf area (*i.e.* by increasing specific leaf area), or by growing faster with existing leaf area (*i.e.* by increasing net assimilation rate [6]).

Many plant species have evolved tolerance to leaf loss [e.g. 5, 7, 8, 9], which indicates that plants have evolved compensatory growth responses. However, relatively little work has been done to study genetic variation in these compensatory growth responses [8]. Furthermore, tolerance can only evolve when there is heritable variation in compensatory mechanisms within populations [1], and thus for the underlying compensatory growth responses. Therefore, in order to estimate the magnitude of adaptation to changing defoliation regimes, estimations of genetic variation in leaf-loss tolerance and associated compensatory growth responses are critical [10].

Plants have to balance between investments in reserves that allow tolerance to disturbances [7, 11] and growth or reproduction. This would suggest a trade-off between tolerance and performance under no disturbance [5]. However, plants can also tolerate defoliation without investing in reserves: by increased photosynthetic activity due to less self-shading, or by higher stomatal conductance due to changed root-shoot ratio [7, 8]. If this is the case, growth under undamaged conditions and tolerance would be expected to be uncorrelated or even positively correlated. The trade-off between growth and tolerance is believed to be a significant factor in determining species habitat adaptation [12]. If tolerance and performance under unstressed conditions are negatively correlated, this could explain the maintenance of genetic diversity in populations with varying levels of disturbance, while a positive genetic correlation is expected to favour superior genotypes and increase variation in life histories among individuals. So far very little is known about the level of within-population genetic correlations between tolerance and performance under unstressed conditions.

Many studies have evaluated genetic variation in performance in short-lived species (mostly annuals and bi-annuals), and some genetic variation in tolerance and genetic correlations between performance and tolerance to leaf-loss [13]. However, for long-lived woody plant species much less is known about these issues [14]. Haukioja & Koricheva [15] argue that tolerance to defoliation might be just as important for long-lived species as it is for short-lived species, but this has not been empirically tested. Defoliation tolerance might be especially relevant for understorey species because shade tolerance is often associated with storage of reserves that allow recovery after damage [12, 16]. More information on the existence of genetic variation in performance, tolerance and genetic correlations between these two, would increase our understanding of the adaptive ability of long-lived plant populations to environmental changes.

In this study we analyzed the extent to which growth and tolerance to defoliation are heritable and if these two variables are genetically correlated. We did this for the long-lived, shade tolerant, tropical understorey palm *Chamaedorea elegans*. Leaf loss due to herbivory and physical damage is high and an important factor limiting the performance of this species [17, 18]. *C. elegans* has been shown to compensate for leaf loss, by changing net assimilation rate (NAR) and allocation of biomass to leaf mass [6]. Furthermore, the leaves of this species are a non-timber forest product, and populations of this species are under pressure due to increased harvesting activities [19].

Specifically, we answered the following questions for our study population:

1. Is there evidence of genetic variation in plant growth and related parameters?
2. Is there evidence of genetic variation in tolerance to defoliation (in terms of growth rate), compensatory growth, and compensatory growth responses (*i.e*. changes in net assimilation rate (NAR), specific leaf area (SLA) and biomass allocation to leaves)?
3. Are growth rate and tolerance to defoliation genetically correlated?

To answer these questions, we performed a greenhouse experiment with seedlings in which a defoliation treatment was applied. We choose to use seedlings because (1) tropical forest seedlings are strongly affected by damage from falling debris and herbivory [16] (2) growing seedlings from collected seeds of mother plants ensured that seedlings were half-sibs (3) using seedlings allowed to increase sample size and obtain results within 1.5 years. We estimated genetic variation in growth parameters, tolerance (in terms of growth), compensatory growth and compensatory growth responses. We used an iterative growth model [6, 20] to estimate NAR, SLA changes, and biomass allocation, which we used to calculate compensation. Furthermore, we analyzed the extent to which tolerance to defoliation and growth rate were related.

## Materials and methods

### Species and site of seed collection

The experiment was performed with the forest understorey palm species *Chamaedorea elegans* Mart, which naturally occurs in rainforest in Mexico, Guatemala, and Belize [21]. It is single stemmed, produces a single cluster of leaves and is dioecious. It naturally occurs mostly on karstic outcrops. Herbivory and falling canopy debris are both major causes of leaf loss in this species [6, 17]. Furthermore, leaves are harvested as a Non-Timber Forest Product (NTFP) for use in the floral industry, causing many populations to be under pressure [19, 22].

Seeds of *C. elegans* were collected from a natural population in south-eastern Mexico in the state of Chiapas. In October 2012, close to the Chajul Biological Station (16°06’ N, 90°56’ W), we set up a 0.7 ha plot, covering the majority of the karstic outcrop where the population was clustered. In this plot, we mapped and tagged all 830 undisturbed individuals with a stem length of >10 cm. From all female fruiting individuals (175 individuals in Nov-Dec 2012) within this plot seeds were collected. In addition, to assure a sufficiently large sample size, seeds were collected from 32 individuals in an 0.1 ha area connected to the main plot in which individuals with a stem length <10 cm were mapped and tagged for a similar experiment in 1997 (using the same methods as in our experiment [17]). In total 3009 seeds from 207 different mother plants were collected, with number of seeds per mother plant ranging from one to 95 seeds. Seeds were cleaned (mesocarp was removed), air-dried and weighed, and they were kept in zip-lock bags that allowed some gas exchange.

### Experimental setup

The experiment was conducted at the Unifarm experimental facilities of Wageningen University, the Netherlands. Seeds were germinated in a growth chamber and later moved to greenhouse. The experiment started for each seedling six months after germination (6 months is an age at which *C. elegans* seedlings growing under the conditions of this experiment have been depleted, S1 File). Plant size was measured non-destructively at the start of the experiment, and a 2/3 defoliation treatment was applied to half of the individuals from each family. The defoliation treatment was repeated every 8 weeks, up to the age of 12.5 months, when plant biomass and other parameters were measured destructively. Details on measurements are provided below in the Data collection and curation section. The timeframe of the experiment (*i.e.* 6.5 months) is similar to other experiments studying tolerance-performance trade-offs in seedlings of long-lived species [11, 16], and was considered to likely be long-enough to reveal differences in allocation of assimilates to storage rather than growth (one of the main mechanisms explaining growth-tolerance trade-offs) [16, 23].

### Germination and greenhouse conditions

In January 2013, seeds were planted at approximately 0.5 cm depth in large trays filled with potting soil. The tray was placed in a growth chamber, where the temperature was kept constant at 30°C day and night, air humidity at 90%. Germination of individual seeds was recorded two times a week. One and a half weeks after emergence, seedlings were transplanted into small pots of 8.5 x 8.5 x 9.5 cm (l x w x h), filled with low nutrient soil (40% peat moss peat, 20% Nordic fraction 2, 20% Baltic peat agent, 20% normal garden peat, 1% pg mix, 0.2% Micromax) and moved to a greenhouse where they were placed in a cage covered with 75% shade cloth to allow for adjustment to changed climatic conditions. After one week, they were moved to a table with flood system allowing a nutrient solution to be absorbed from below into the pots (pH 5.0, EC 0.8, NPK ratio 12-14-24). Seedlings stayed on the table with flood system for the duration of the experiment (see the Experimental setup section below). To simulate forest conditions, temperature in the greenhouse was kept at a minimum day/night temperature of 24/22°C, air humidity at 80%, day length was reduced to a maximum of twelve hours using automatically closing black screens. Light levels were in summer months reduced using (depending on the month) either 25% or 50% shade cloth, such that plants received approximately 2 mol per day, which is the average light intensity in the forest understorey at the site where seeds were collected [24]. Monthly target shade levels were based on the 10-year monthly average light intensities recorded at the location of the greenhouses.

### Experimental design and treatment

The experiment was laid out as a randomized block design with six blocks. To this end, the table was divided into six equal parts lengthwise to create the blocks. Seedlings from the same mother (half-sib families) were randomly distributed over the blocks and over position within the block. Because families differed in number of seedlings, sometimes a family was only present in one block (this was the case for families with only one seedling), and sometimes in all six (which was the case for families with at least six seedlings).

To assign the seedlings to control or defoliation treatments, we ranked all plants in a family according to age (*i.e*. date of emergence). We then randomly assigned a treatment (*i.e.* defoliation or control) to the oldest one, giving the other treatment to the second oldest plant and alternating in this way across the age hierarchy. Of all seedlings that were assigned to the defoliation treatment, two out of every three leaflets were cut off at six months of age. This treatment was repeated (for newly produced leaves) every eight weeks.

### Data collection and curation

At six months of age, we measured seedling stem length and diameter. In addition, we measured leaf width, lamina length, rachis length, rachis diameter, leaflet width, and number of leaflets of all leaves, as well as the length of unopened leaf. With this information, seedling biomass (per plant part) and leaf area of the seedlings of six months of age were estimated using an allometric model, that we constructed based on data of a destructive harvest of extra seedlings of six months of age from the same experimental conditions (see S1 File for details).

Surviving seedlings were destructively harvested at 12.5 months of age (1387 in total). Plants were checked for natural abscissions (which can easily be detected by the structure of the plant), but no natural abscissions were detected. Roots were carefully washed to remove all soil particles. Leaf area was measured of the second fully developed leaf (counting from the apex), using a leaf area meter (LiCor LI3100 Lincoln NE, USA). Roots, stem, rachis, undeveloped leaves, lamina of non-defoliated leaves and lamina of defoliated leaves were separated, and dried in a stove at 70°C for at least 72 hours, after which dry mass per plant part was determined.

Measured weights and leaf area were checked for mistakes. Mistakes included incomplete defoliation treatment, no separation of non-defoliated leaf mass at harvest, no defoliation of new leaves at harvest, and unrealistic values. Unrealistic values were defined as deviations of more than a factor of ten from the mean observed relative value compared to other plant parts (*e.g.* from the leaf mass/stem mass ratio). A total of 88 plants were excluded from further analysis. From the included individuals, we selected only those that belonged to families (*i.e.* were obtained from a mother palm) that contained at least 12 individuals. The selection reduced the initial number of 207 families sampled in the field to 47 families included in the analyses. Analyses were conducted on a total of 731 seedlings.

### Estimation NAR, biomass allocation to leaves, changes in SLA and RGR

To estimate growth and several growth-related variables (net assimilation rate (NAR), fraction of newly assimilated mass that is allocated to lamina growth (f_lam_), fraction in daily change in mean specific leaf area (γ) and relative growth rate (RGR)), we used an iterative growth model following the method of Anten & Ackerly [20]. This method of growth analysis allows more exact estimations of growth variables than either the classic or functional approaches of growth analysis [25] when a plant experiences repeated defoliation because it includes timing of leaf loss [20]. Input for this model is biomass, leaf mass, and leaf area at the beginning and end of the experiment, and leaf loss (mass and area, and time of removal) during the experiment. We, however, did not measure leaf loss directly but assumed this to be two third of existing leaf mass (*i.e.*, our defoliation treatment entailed removing two out of every three leaflets). To allow for this, we adjusted the Anten & Ackerly [20] model. A more detailed description of these methods is provided in S2 File.

### Estimation of tolerance and compensatory responses

Tolerance and compensatory growth are both measures of plant performance under defoliation stress, compared to performance of control (non-defoliated) plants. Tolerance, the difference in fitness (or growth in our case) between individuals under defoliation stress and non-defoliated individuals [1], is the measure most widely used to make such comparisons, but it does not take into account the amount of leaf area that was removed. Compensation, the fraction of the potential loss in growth due to leaf loss that is mitigated through compensatory mechanisms, does take lost leaf area into account and some methods allow for including the time of removal as well [20]. This more functional approach allows for estimation of the underlying growth parameters (changes in NAR, SLA and biomass allocation). We analysed both, because growth tolerance is a more common measure, but compensation gives more insight in the underlying mechanisms.

To be able to estimate genetic variation in tolerance and compensation, information on differences in tolerance within families, and therefore per individual is required. In order to be able to calculate tolerance and compensation per individual, each individual in the defoliation treatment was paired with a family member from the control treatment, based on rank order of estimated biomass at six months of age (*i.e.* seedling age at the beginning of the experiment). Pairing is a standard procedure in growth analysis [1]. Using the values of the coupled control individual, tolerance in growth rate was calculated as T = (G_D_-G_C_)/G_D_ in which T indicates tolerance, G growth, and the subscript D and C the defoliation- and control treatment respectively. For tolerance in RGR, RGR values were obtained with the iterative growth model. For tolerance in biomass growth, we calculated biomass change between 6 months and 12 months of age, for which the values were obtained from direct measurements. We excluded leaf mass in this calculation.

We estimated compensatory growth per individual using the approach of Anten, et al. [6]. We used the coupled control family members as a null-model to be able to estimate growth rate of a hypothetical, non-compensating individual. Using the start-biomass of the defoliated individual, but the growth parameters (NAR, f_lam_, γ) of the control individual, we calculated biomass growth rate and RGR based on the iterative growth model, for both the control and defoliation treatment. Compensation was then calculated as *Compensation*= 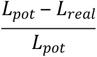 in which L_pot_=C0-D0 and L_real_=C0-D. L_pot_ (the potential reduction in growth) is therefore calculated as the growth of a control individual with the null-model growth parameters (C0), minus growth of a defoliated individual with the same null-model growth parameters (D0). L_real_ (the realized reduction in growth) is calculated as C0 minus the actually realized growth of the defoliated individual (D).

### Statistical analysis

To estimate genetic variation in growth parameters (NAR, f_lam_ and γ), variables of biomass growth (without leaf mass) and RGR, and for tolerance and compensation, we constructed mixed effect models, in which (half-sib) family (F) was included as random factor. Seed weight (s) was included as fixed effect when its effect was significant, to correct for potential maternal effects. The resulting models were *y_ij_ = μ + S_j_ + F_i_ + e_ij_* and *y_ij_ = μ + F_i_ + e_ij_* with F_i_ ~ N(0,σ_F_^2^) and e_ij_ ~ N(0,σ^2^). From the among-family variance component 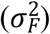 and the residual variance component (σ^2^) narrow sense heritability was estimated as 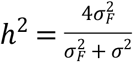. Because mother plants were randomly pollinated, families were considered to be half-sibs in this estimation [26]. Estimates for plants that were part of the defoliation treatment were calculated separately.

To analyze genetic variation in response to defoliation, we constructed mixed effect models for all estimated growth parameters in which treatment (T) was included as a fixed effect, family as a random effect, as was the interaction term between treatment and family. A relatively large interaction term between defoliation treatment and family in the models of biomass growth or RGR, is an indication of genetic variation in tolerance [e.g. 27]. Likewise, a relatively large interaction term between treatment and family in the mixed models for the growth parameters NAR, f_lam_ and γ, are indications of genetic variation in compensatory traits. When visual inspection of the data suggested more complex variance structures, these were modeled as well, and the best model was selected based on Akaike (AIC) criteria. The best model was for all tested variables the model in which separate within group variance components were estimated per treatment, which is *y_ijk_ = μ + T_j_ + s_k_ + F_i_ + F × T_ij_ + e_ijk_* with F_i_ ~ N(0,σ_F_^2^), F × T_ij_~ N(0,σ_FxT_^2^) and e_ijk_ ~N(0,σ_j_^2^). Mixed effect models were analyzed in Genstat [28], all other analyses were performed in R [29].

## Results

### Genetic variation in growth parameters

We found large variation among different families in biomass growth and RGR (Fig 1). We determined within and among family variance components for biomass growth rate, RGR, and the growth parameters NAR, biomass allocation (f_lam_), and SLA change (γ) that were estimated by the iterative growth model (Table 1). Based on the gathered variance components, we estimated narrow-sense heritability of growth rate to be relatively large for non-defoliated plants, and only slightly lower for plants that were subjected to defoliation (*h*^2^ values for biomass growth and RGR ranged from 0.41 to 0.46 for control plants and from 0.32 to 0.35 for defoliated plants, Table 1). Surprisingly, estimations of heritability of the growth parameters NAR, f_lam_, and γ, were much lower, especially for the control individuals (Table 1).

**Fig. 1.**
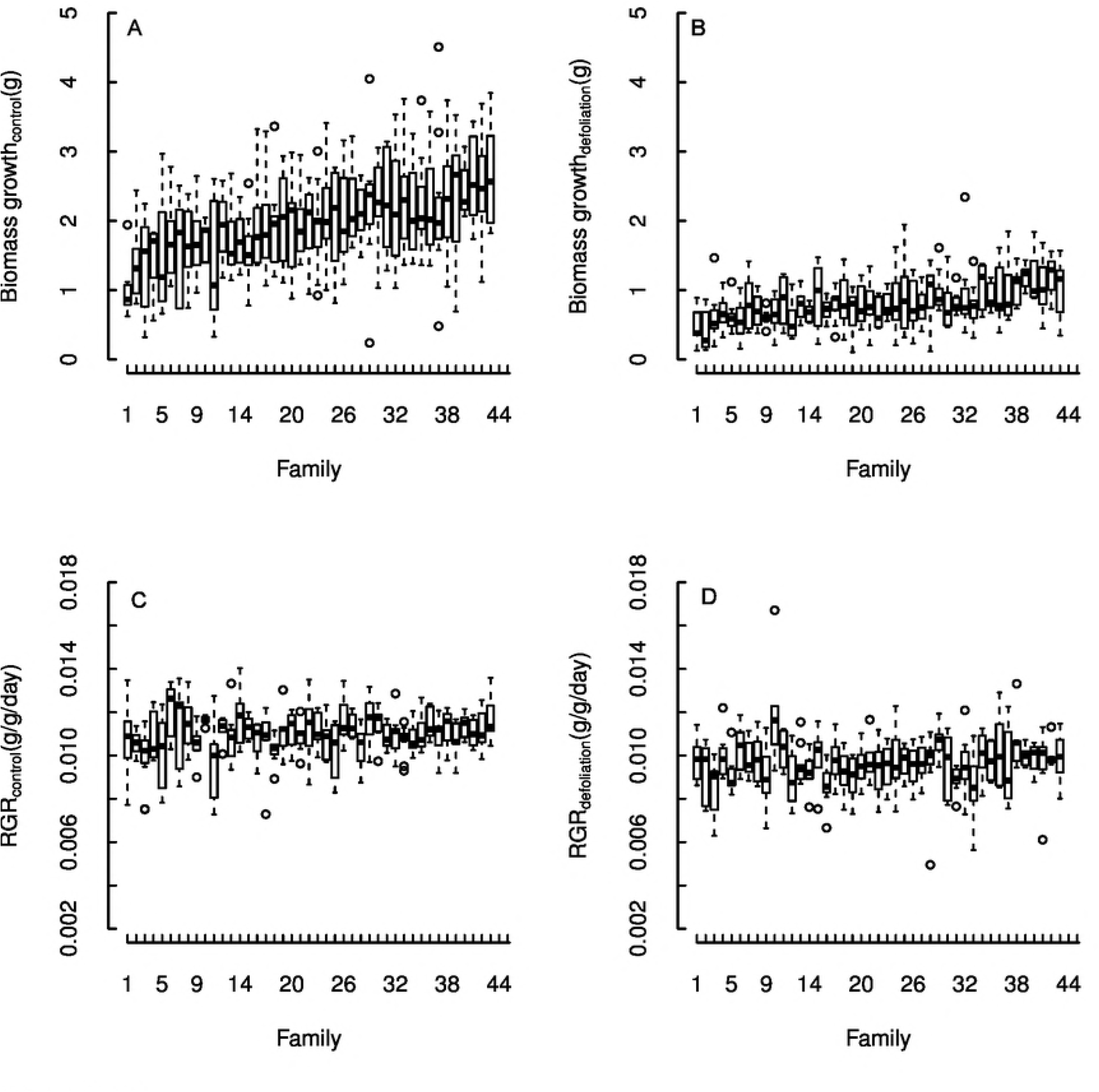
Boxplots of biomass growth and RGR for control and defoliated seedlings of 47 families of *Chamaedorea elegans* from a Mexican rainforest. Boxes are the interquartile range (IQR), black lines in the middle of boxes are medians, whiskers are the extreme data point with 1.5 x IQR. Families are ranked by increasing order of mean biomass growth. The changing rank of families between treatments is a first indication that families that grow relatively fast without the stress of defoliation do not necessarily grow relatively fast when they suffer leaf loss. The changes in rank between biomass growth rate and RGR indicate that families that grew fast in absolute terms did not necessarily grow fast in relative terms.

**Table 1.**
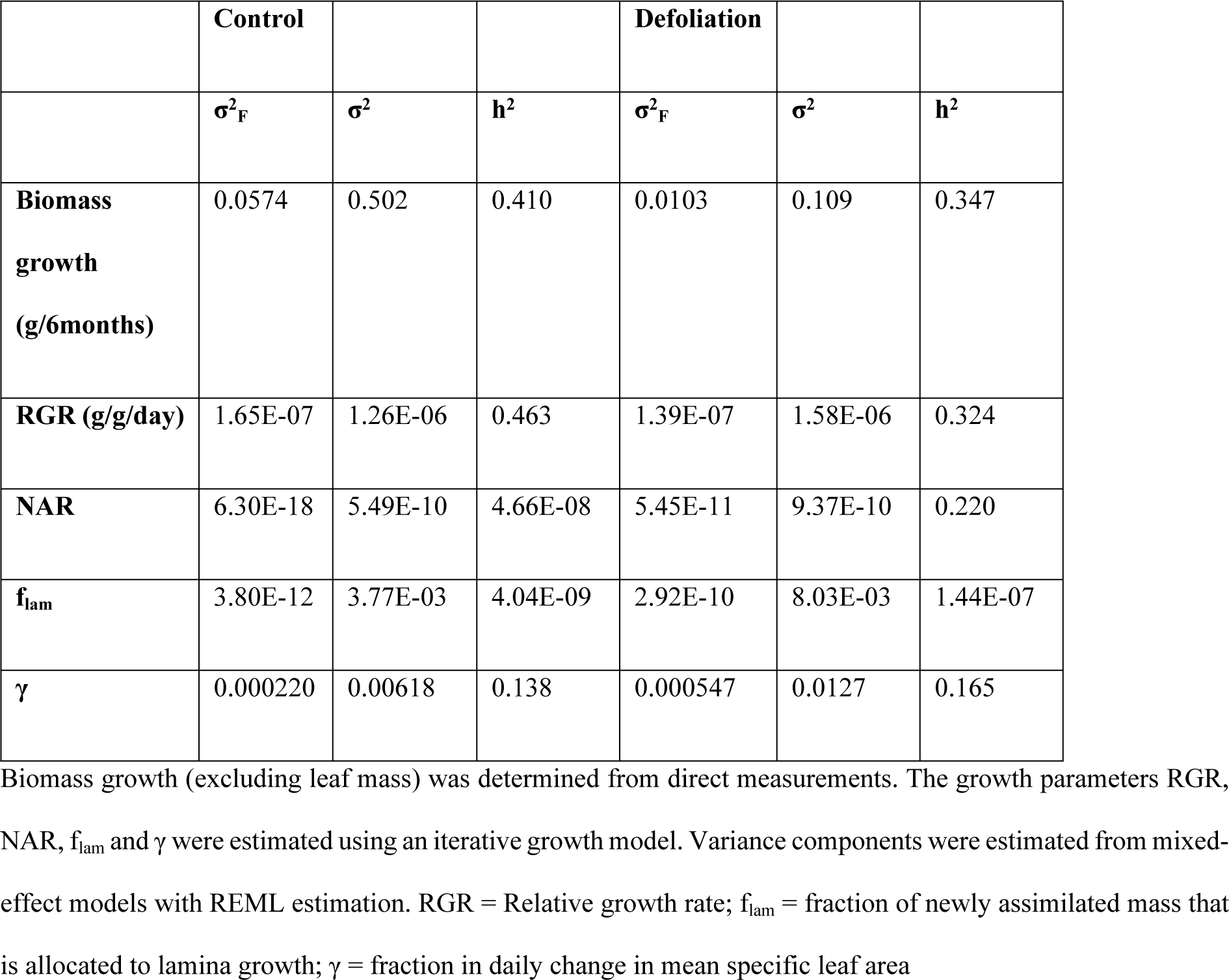
Estimated within- and among-family variance components and narrow-sense heritability (*h*^2^) for several growth parameters for a population of the understorey palm *Chamaedorea elegans*, for which seedlings were subjected to defoliation in a greenhouse.

### Genetic variation in tolerance, compensation, and compensatory traits

We compared family mean control and defoliation treatment values of all growth parameters (Fig 2). Family mean biomass growth rate was as expected, lower in the defoliation treatment for all families and for RGR in almost all families. However, all family mean values of NAR and biomass allocation, and almost all family mean values of SLA change, were higher in the defoliation treatment than in the control treatment. Therefore, all families clearly showed compensatory responses to leaf loss by increasing their NAR and SLA, and changing their biomass allocation.

**Fig. 2.**
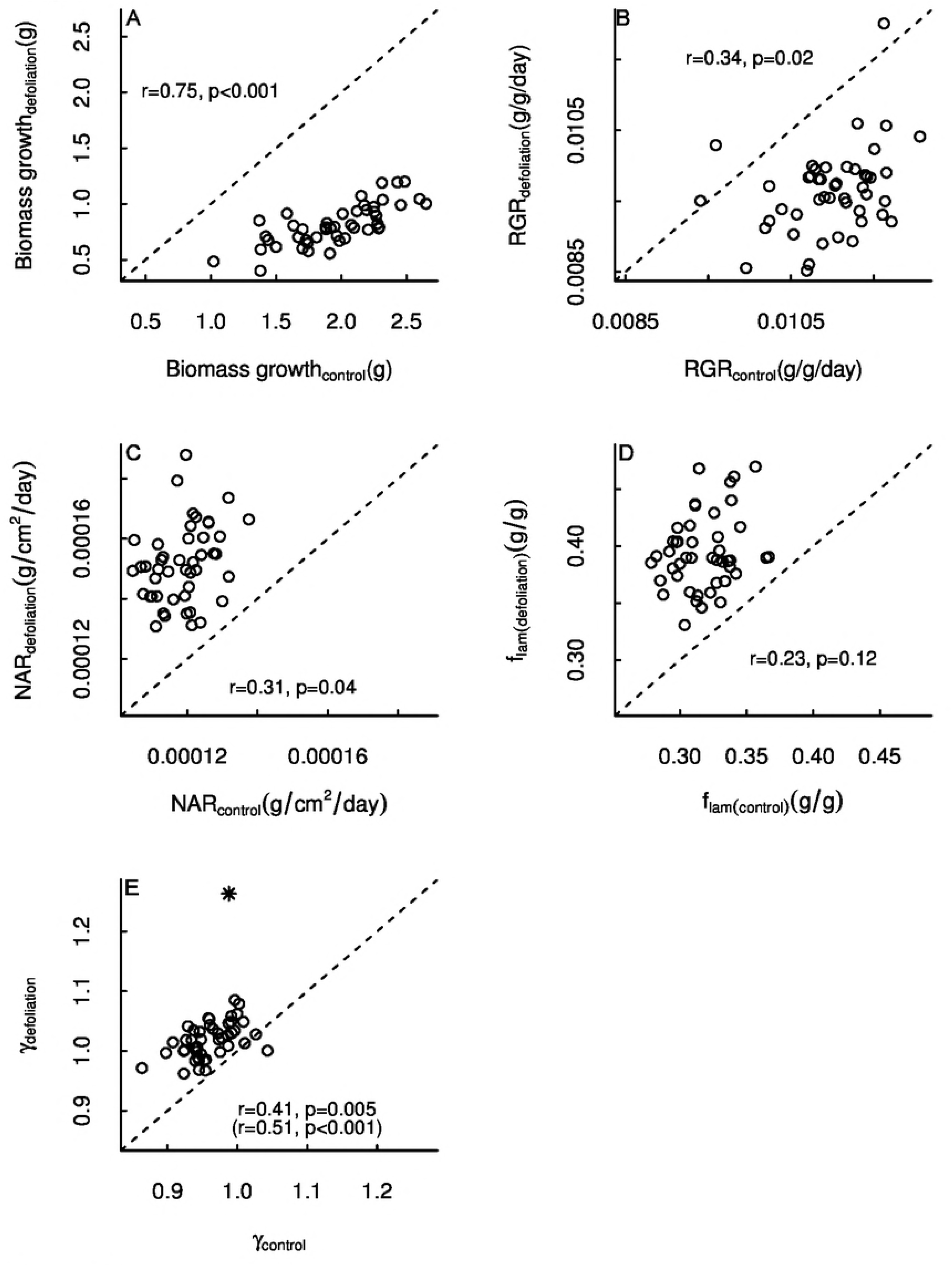
Comparison of control and defoliation treatment family means of several growth parameters for seedlings of the understorey palm *Chamaedorea elegans*. Biomass growth was determined from direct measurements, the other parameters were all estimated using an iterative growth model. The dashed line indicates a 1-to-1 relationship. Pearson correlation coefficients and associated p-values are shown. The asterisk in panel (e) is an outlier data point; correlation coefficient and p-value without this data point are shown in between brackets.

We tested whether families responded differently to defoliation, and therefore whether there was genetic variation in response to defoliation, with a mixed effect model in which we included the random interaction between treatment and family. This model yielded only relatively small variance components for the interaction between treatment and family for all evaluated parameters (Table 2). This suggests that families do not respond significantly different to leaf loss in terms of biomass growth, RGR, NAR, allocation to leaf mass nor SLA changes. Therefore, while families compensate strongly for leaf loss, we did not find evidence for strong within-population genetic variation in this response.

**Table 2.**
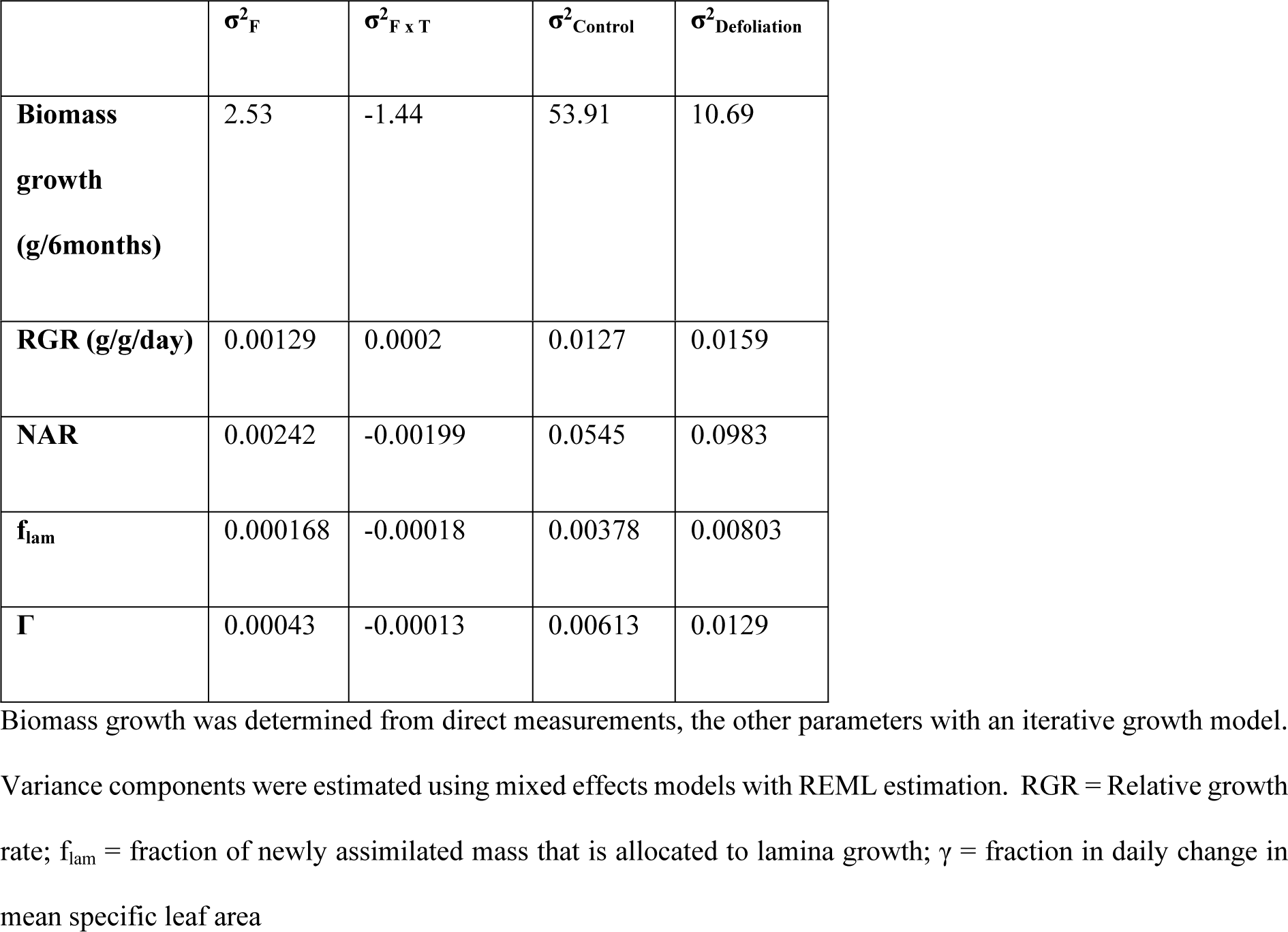
Estimated family, family*treatment and residual variance components for several growth parameters, estimated from a greenhouse experiment that was performed with seedlings for which the seeds came from a small (0.7ha) Mexican population of the understorey palm *Chamaedorea elegans*.

To estimate genetic variation in tolerance and compensation itself, we paired defoliation treatment individuals with control individuals from within the same family. By doing this, we were obtaining replicated estimates of tolerance and compensation and could therefore estimate the heritability of these parameters. Even though we found large variation between family mean values of tolerance and compensation (*e.g.* family mean compensation in biomass growth ranged from 0.16 to 1.03, *i.e.*, 16 – ~100% of potential loss being mitigated), within-family variance was much larger. Therefore, estimations of heritability of tolerance and compensation were low (the highest estimated heritability was for compensation in biomass growth, which was only 0.01, Table 3).

**Table 3.** Estimated within and among family variance components and heritability of tolerance to defoliation, and compensation after repeated defoliation events in a greenhouse experiment, performed seedlings of the understorey palm *Chamaedorea elegans*. To be able to estimate tolerance and compensation, individuals from the defoliation treatment were coupled to individuals from the control treatment based on their estimated biomass at the start of the experiment. Compensation was calculated by using an iterative growth model that allowed estimation of a hypothetical non-compensating individual.

|  |  |  |  |  |
| --- | --- | --- | --- | --- |
|  | **Tolerance** |  |  | **Compensation** |

| | $\sigma^2_{\text{Family}}$ | $\sigma^2$ | $h^2$ | $\sigma^2_{\text{Family}}$ | $\sigma^2$ | $h^2$ |
| --- | --- | --- | --- | --- | --- | --- |
| **Biomass growth (g/6months)** | 0.00636 | 2.796 | 0.00908 | 0.000559 | 0.1820 | 0.0122 |
| **RGR (g/g/day)** | 1.53E-10 | 6.18E-02 | 9.90E-09 | 1.76E-09 | 5.23E-01 | 1.35E-08 |

Note: RGR = Relative growth rate

### Relation between growth and tolerance

For all growth parameters, there were positive correlations between family mean control values and family mean defoliation treatment values, indicating that growth performance was genetically correlated between treatments (Fig 2). The correlation coefficient for biomass growth was higher (r = 0.75) than those for RGR, NAR and γ (r = 0.34, r = 0.31, and r = 0.41 respectively). Only the estimated positive correlation coefficient of f_lam_ (r = 0.23) was not significant. These results suggest the existence of superior genotypes that grow fast while still being able to tolerate defoliation.

It is possible that even though (to some extent) the same families grew faster in both treatments, the relative reduction in growth rate might have been larger for families that grew fast in the control treatment. If this was the case, there would be a negative relation between tolerance or compensation (both relative measures) and growth rate in the control treatment. To test this we compared family mean values of tolerance and compensation, to family mean values of biomass growth rate and RGR in the control treatment (Fig 3). This did not yield clear evidence for any positive or negative relation between tolerance/compensation and biomass growth/RGR. The only significant correlation that we found was between tolerance and RGR. However, this relationship was heavily pulled by two outlying data points; without these outliers there was no longer a significant correlation. Therefore, we did not find evidence that would suggest costs to tolerance in terms of growth.

**Fig. 3.**
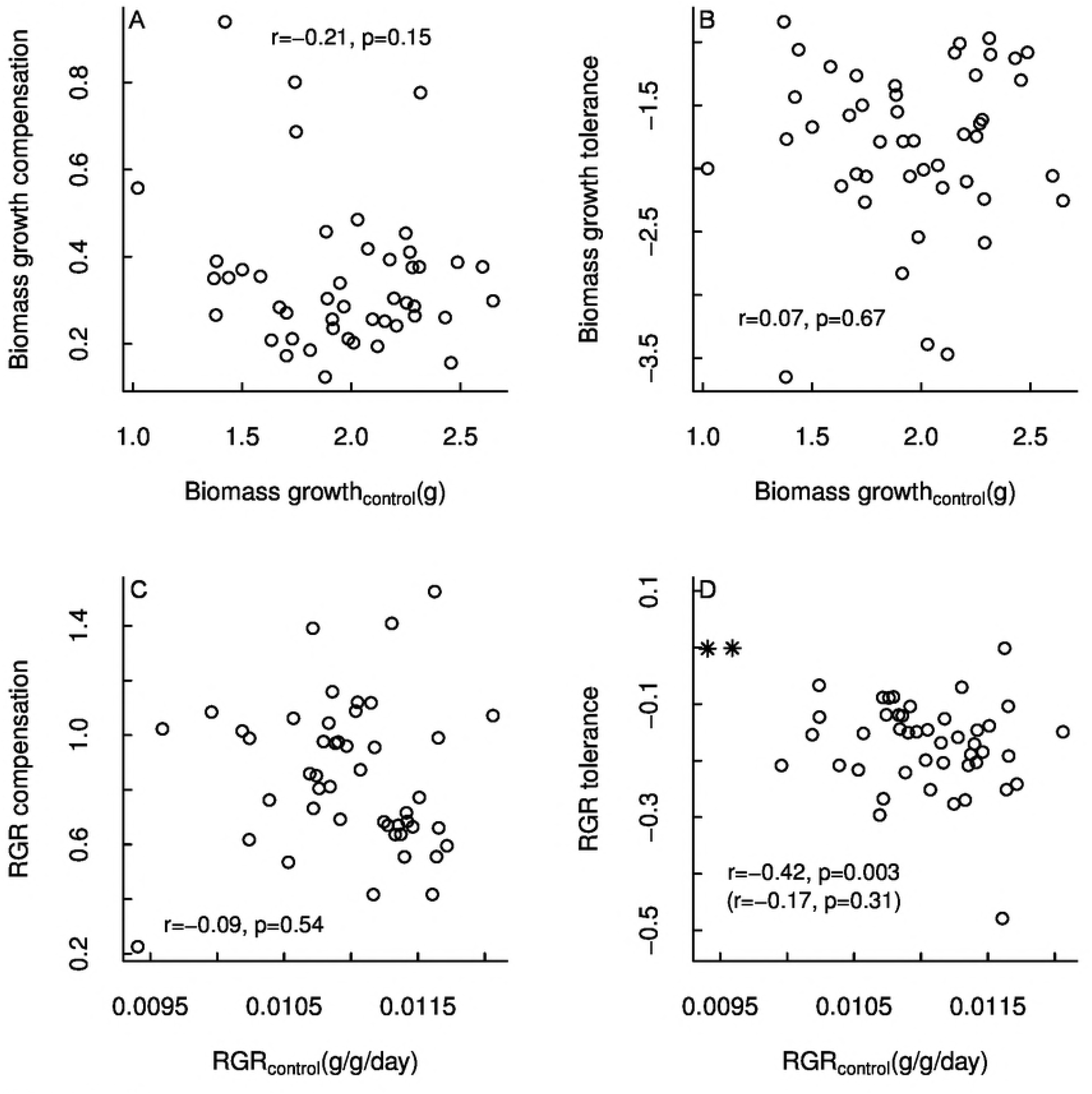
Relationships between family mean compensation (A, C), tolerance (B, D) and family mean growth rate. Data were obtained from 47 half-sib families of seedlings of the understorey palm *Chamaedorea elegans*, in which a defoliation treatment was applied. Compensation, RGR tolerance and RGR were estimated with an iterative growth model that takes into account timing of leaf removal (see methods). Pearson correlation coefficients and associated p-values are provided. The asterisks in panel d are two outlying data points; Pearson correlation coefficient and p-value without these data points are shown in between brackets.

## Discussion

This study showed that genetic variation in tolerance and compensatory responses to defoliation is limited within a population of a long-lived tropical forest species. We also showed that genetic variation in growth potential was much larger than values usually detected for small populations [14, 30]. These results suggest that the studied population might have limited ability to adapt in terms of tolerance to environmental changes that entail leaf loss but does have the ability to adapt to environments that require different growth rates. Furthermore, this is one of the first studies that has analyzed genetic variation in compensatory growth responses to defoliation.

### Heritability of growth potential

We found large within-population genetic variation in growth rate, with estimations of narrow-sense heritability ranging from 0.32 to 0.46. These estimations are higher than the estimations from the few other studies that have been performed with long-lived plant species. For example, in the shade tolerant rainforest tree *Sextonia rubra* heritability ranged from 0.23 to 0.28 for several growth-related traits [30], and between 0.20 and 0.37 in a population of *Populus tremuloides* [14]. The values that we found are especially high considering that the seeds used in this experiment were collected in a very small area (0.7 ha). Furthermore, the high genetic variation that we found is somewhat surprising because inbreeding in *Chamaedorea* species has been estimated to be high in several other Mexican *C. elegans* populations [31]. This suggests that heritability in growth could be higher in understorey palms than in trees, but further research on multiple populations and species is necessary to determine this.

### Compensatory responses and heritability of tolerance to defoliation

We found individuals to compensate strongly for defoliation, by increasing NAR, allocating more biomass to leaf mass, and by increasing SLA, which are similar responses that have been found in other studies [e.g. 32] including one that was also performed with *C. elegans* [albeit with adults, 6]. Mean families values of compensation varied strongly (*e.g.* for biomass growth between 0.16 to 1.03, *i.e.*, the extent of compensation from about 1/8 to full compensation). However, we found only very limited evidence for genetic variation in compensatory responses and tolerance. Genetic variation in tolerance has been found for many species of annual and bi-annual plants (see *e.g*. [1]for a review on this), but, as Stevens, Waller & Lindroth [14] point out, much less is known about the level of genetic variation in tolerance in long-lived species. A reason for this is that resistance (*e.g.* chemical defenses) rather than tolerance has long been seen as a more effective measure for long-lived species to persist under the pressure of herbivory, due to their different life-history traits, such as long-lived leaves [15]. However, as explained by Haukioja & Koricheva [15], tolerance could be just as important for long-lived species as for the short-lived ones, partly because herbivore attacks can never be completely avoided, and plants endure leaf losses due to chronic physical damages. Tolerance could be particularly well developed in understorey species because shade tolerance is often associated with storage of reserves that allow recovery after damage [12, 16, 23] and because understory plants are subjected to falling canopy elements like branches, limbs and complete trees [33]. Studies that have been performed on long-lived plants were all on tree species (in which part of the studies detected genetic variation in tolerance, *e.g*. [14], while others did not, *e.g*. [34]. To our knowledge, genetic variation in tolerance and compensatory responses has not been studied in natural populations of other types of long-lived plant species like lianas, ferns or palms.

### Relation between growth and tolerance

We did not detect a genetic correlation between growth and tolerance or compensation, even though it has been shown that such correlation exists at least at the ecotype level in short-lived plants [32]. Therefore, the strong differences in growth that we detected among families cannot be explained by a growth-tolerance trade-off. In contrast, we found that ‘super-performing’ families that grew relatively fast under undisturbed conditions also grew fast when exposed to defoliation. These types of superior genotypes could play a key role in population resistance when the population is being disturbed by, for example, a storm (and associated increase of falling canopy debris) or herbivore attack. Fast growers have been shown to contribute positively and disproportionately to population growth [35, 36], and our results suggest that such contribution would be maintained under disturbance. However, population growth is not only influenced by the response of individuals to disturbance in terms of growth but also by their survival and ability to maintain seed production under stress. Therefore, it would be very interesting to test if fast growing adult plants have a high survival probability and are better able to maintain seed production when they suffer leaf loss, especially because *Chamaedorea spp*. have been shown to be relatively intolerant to leaf loss in terms of reproduction [6, 22, 37].

A trade-off with defoliation tolerance did not explain why genetic diversity for growth potential was high within the population that we studied. However, it is possible that there are other trade-offs with growth than the one with defoliation tolerance such as genotype x environment trade-offs (*i.e*. G x E interactions). Our study site is characterized by persistent spatial heterogeneity in environmental conditions [38]. Possibly, genotypes that grow fast in certain environmental conditions, like the greenhouse conditions in this experiment, are not the ones that would grow fast in other environments that are, for example, nutrient poor. However, it is hard to estimate how likely this is, as G x E interactions have hardly been studied in long-lived plant species, in particular, those that occur in tropical forests.

The current study was performed with seedlings. Possibly, our estimations of genetic variation in tolerance and compensatory growth responses could be different if the experiment had been performed with adult plants. Larger reserve storage in adult plants may lead to higher tolerance to defoliation compared to seedlings. However, compensatory responses were strong in our experiment and comparable to those reported for adults of the same species [6], suggesting that if genetic variation in these responses would be strong in our study population, this would have been expressed in our experiment.

### Implications

The low genetic variation in compensatory responses and tolerance that we found, could have consequences for the adaptive potential of populations to environmental changes [10]. If the frequency and magnitude of leaf loss in a population persistently increases (*e.g.* due to an increase of storm frequencies, which is predicted in several climate change scenarios [39], or due to the introduction of an invasive herbivore [40]), populations with limited genetic variation in tolerance to defoliation might not be able to respond and adapt to such selective pressures. On the contrary, the high genetic variation that we estimated for growth potential, might increase the adaptability of populations if pressure for light competition changes. This could, for example, happen if canopy dynamics change due to differences in storm frequencies, or because of the introduction of a new faster-growing, light-demanding, understorey species. In this case, genotypes that allow high growth might be selected for.

In the above context, it is critical to obtain accurate information on genetic variation in quantitative traits present in populations in order to be able to evaluate what the effect of environmental change will be on populations [10]. Especially information on genetic variation in traits that are directly linked to individual vital rates is essential to be able to link evolutionary and demographic processes [41]. However, at this point, surprisingly little is known about this for tropical forest species. Therefore, we strongly recommend more studies that evaluate the amount of within-population genetic variation causing differences in vital rates, and the consequences of this variation for the adaptive potential of populations to changing environments.

Strong genetic variation in growth rate as we found in this study, can also have implications for management practices. The existence of superior individuals that grow faster while still being able to strongly compensate for leaf loss offers opportunities for increased production by artificial selection. These individuals can be used when a species is commercialized, especially when this is for its leaves. In the case of *C. elegans*, leaves are harvested as a non-timber forest product (NTFP) for the floral industry, and are increasingly being planted in secondary forests for enrichment or in intercropping systems with species that provide shade [42]. This study shows that it might be beneficial to select seeds from individuals that have high growth rates, which can be easily identified for this species [35]. We believe that there are many more long-lived tropical forest species for which it could be valuable to explore this potential.

## Acknowledgements

We thank Unifarm personnel for technical support and logistic help in the design and execution of the experiment, and Albert van der Veen, Mitchel Jansen, and Ruud Grootens for help with data collection during their internship. We also like to thank the Gideon Terburg, Stefan van Meijeren, and various volunteers that helped with collection of data, in particular Laura Mackenbach. Chris Maliepaard is thanked for his advice on the setup of the experiment, and Arthur Spruit of Aardam Plants for sharing his knowledge on seed germination and best seedling growth condition of *C. elegans*. We thank the Chajul Field Station for facilities provided to conduct this research.

**Supporting Information**

**S1 File. Allometric model.** Details on methods of the construction of an allometric model for estimation of biomass per plant part of seedlings of 6 months of age

**S2 File. Iterative growth model.** Details on methods of the construction and adaptation of an iterative growth model for estimation of daily individual seedling NAR, flam and γ

## References

1. Strauss SY, Agrawal AA. The ecology and evolution of plant tolerance to herbivory. Trends Ecol Evol. 1999;14(5):179–85. Epub 1999/05/14. PubMed PMID: 10322530.

2. Núñez-Farfán J, Fornoni J, Valverde PL. The evolution of resistance and tolerance to herbivores. Annual Review of Ecology, Evolution, and Systematics. 2007:541–66.

3. Belsky AJ, Carson WP, Jensen CL, Fox GA. Overcompensation by plants: herbivore optimization or red herring? Evolutionary Ecology. 1993;7(1):109–21.

4. Agrawal AA. Overcompensation of plants in response to herbivory and the by-product benefits of mutualism. Trends in plant science. 2000;5(7):309–13.

5. Stowe KA, Marquis RJ, Hochwender CG, Simms EL. The evolutionary ecology of tolerance to consumer damage. Annual Review of Ecology and Systematics. 2000:565–95.

6. Anten NPR, Martínez-Ramos M, Ackerly DD. Defoliation and growth in an understorey palm: Quatifying the contributions of compensatory responses. Ecology. 2003;84:2905–18. PubMed PMID: 220.

7. Tiffin P. Mechanisms of tolerance to herbivore damage: what do we know? Evolutionary Ecology. 2000;14(4-6):523–36.

8. Anten NPR, Pierik R. Moving resources away from the herbivore: regulation and adaptive significance. New Phytologist. 2010;188(3):643–5.

9. Fornoni J. Ecological and evolutionary implications of plant tolerance to herbivory. Functional Ecology. 2011;25(2):399–407. doi: 10.1111/j.1365-2435.2010.01805.x.

10. Lande R, Shannon S. The role of genetic variation in adaptation and population persistence in a changing environment. Evolution. 1996;50(1):434–7.

11. O’Brien MJ, Burslem DF, Caduff A, Tay J, Hector A. Contrasting nonstructural carbohydrate dynamics of tropical tree seedlings under water deficit and variability. New Phytologist. 2015;205(3):1083–94.

12. Kobe RK. Carbohydrate allocation to storage as a basis of interspecific variation in sapling survivorship and growth. Oikos. 1997:226–33.

13. Geber MA, Griffen LR. Inheritance and natural selection on functional traits. International Journal of Plant Sciences. 2003;164(S3):S21–S42.

14. Stevens MT, Waller DM, Lindroth RL. Resistance and tolerance in Populus tremuloides: genetic variation, costs, and environmental dependency. Evolutionary Ecology. 2007;21(6):829–47.

15. Haukioja E, Koricheva J. Tolerance to herbivory in woody vs. herbaceous plants. Evolutionary Ecology. 2000;14(4-6):551–62.

16. Myers JA, Kitajima K. Carbohydrate storage enhances seedling shade and stress tolerance in a neotropical forest. Journal of Ecology. 2007;95(2):383–95.

17. Martínez-Ramos M, Anten NPR, Ackerly DD. Defoliation and ENSO effects on vital rates of an understorey tropical rain forest palm. Journal of Ecology. 2009;97:1050–61. doi: 10.1111/j.1365-2745.2009.01531.x. PubMed PMID: 97.

18. Valverde T, Hernandez-Apolinar M, Mendoza-Amaro S. Effect of leaf harvesting on the demography of the tropical palm Chamaedorea elegans in South-Eastern Mexico. Journal of Sustainable Forestry. 2006;23(1):85–105.

19. Reining CC, Heinzman RM, Madrid MC, Lopez S, Solorzano A. Non-timber forest products of the Maya Biosphere Reserve, Petén, Guatemala. Non-timber forest products of the Maya Biosphere Reserve, Peten, Guatemala. 1992.

20. Anten NPR, Ackerly DD. A new method of growth analysis for plants that experience periodic losses of leaf mass. Functional Ecology. 2001;15(6):804–11. PubMed PMID: 267.

21. Hodel DR. Chamaedorea palms, the species and their cultivation. Lawrence, Kansas, USA: Allen press; 1992.

22. Hernández-Barrios JC, Anten NPR, Ackerly DD, Martínez-Ramos M. Defoliation and gender effects on fitness components in three congeneric and sympatric understorey palms. Journal of Ecology. 2012;100(6):1544–56. doi: 10.1111/j.1365-2745.2012.02011.x.

23. Poorter L, Kitajima K. Carbohydrate storage and light requirements of tropical moist and dry forest tree species. Ecology. 2007;88(4):1000–11.

24. Anten NPR, Ackerly DD. Canopy-level photosynthetic compensation after defoliation in a tropical understorey palm. Functional Ecology. 2001;15(2):252–62. doi: 10.1046/j.1365-2435.2001.00517.x.

25. Poorter H. Plant growth analysis: towards a synthesis of the classical and the functional approach. Physiologia Plantarum. 1989;75(2):237–44.

26. Falconer DS. Introduction to quantitative genetics. 4th ed., &lt repr.&gt ed. Mackay TFC, editor: Harlow : Longman; 1997.

27. Agrawal AA, Strauss SY, Stout MJ. Costs of induced responses and tolerance to herbivory in male and female fitness components of wild radish. Evolution. 1999:1093–104.

28. International V. GenStat for Windows 14th Edition Hemel Hempstead, UK: VSN International; 2011.

29. R Development Core Team. R: A language and environment for statistical computing. Vienna, Austria: R Foundation for Statistical Computing; 2014.

30. Bonal D, Scotti I, Calvo-Vialettes L, Scotti-Saintagne C, Citterio M, Degen B. Genetic variation for growth, morphological, and physiological traits in a wild population of the Neotropical shadetolerant rainforest tree Sextonia rubra (Mez) van der Werff (Lauraceae). Tree genetics and genomes. 2010;6(2):319–29. PubMed PMID: 95.

31. Cibrián-Jaramillo A, Hahn WJ, Desalle R. Development of microsatellite markers of the Mexican understorey palm Chamaedorea elegans, cross-species genotyping, and amplification in congeners. Molecular Ecology Resources. 2008;8(2):322–4. PubMed PMID: 263.

32. Camargo ID, Tapia-López R, Núñez-Farfán J. Ecotypic variation in growth responses to simulated herbivory: trade-off between maximum relative growth rate and tolerance to defoliation in an annual plant. AoB plants. 2015;7:plv015.

33. Martinez-Ramos M, Alvarez-Buylla E, Sarukhan J, Pinero D. Treefall age determination and gap dynamics in a tropical forest. The Journal of Ecology. 1988:700–16. doi: 10.2307/2260568.

34. Axelsson EP, Hjältén J. Tolerance and growth responses of populus hybrids and their genetically modified varieties to simulated leaf damage and harvest. Forest Ecology and Management. 2012;276:217–23.

35. Jansen M, Zuidema PA, Anten NPR, Martínez-Ramos M. Strong persistent growth differences govern individual performance and population dynamics in a tropical forest understorey palm. Jounal of Ecology. 2012;100 (5):1224–32. doi: 10.1111/j.1365-2745.2012.02001.x. PubMed PMID: 242.

36. Zuidema PA, Brienen RJW, During HJ. Do Persistently Fast-Growing Juveniles Contribute Disproportionately to Population Growth? A New Analysis Tool for Matrix Models and Its Application to Rainforest Trees. American naturalist. 2009;174(5):709–19. PubMed PMID: 123.

37. Endress BA, Gorchov DL, Noble RB. Non-timber forest product extraction: effects of harvest and browsing on an understory palm. Ecological applications. 2004;14:1139–53. PubMed PMID: 158.

38. Jansen M, Anten NPR, Bongers F, Martínez-Ramos M, Gavito ME, Zuidema PA. Explaining long-term inter-individual growth variation in plant populations: persistence of abiotic factors matters. Oecologia. 2017;185(4):663–74.

39. Dale VH, Joyce LA, McNulty S, Neilson RP, Ayres MP, Flannigan MD, et al. Climate change and forest disturbances: climate change can affect forests by altering the frequency, intensity, duration, and timing of fire, drought, introduced species, insect and pathogen outbreaks, hurricanes, windstorms, ice storms, or landslides. BioScience. 2001;51(9):723–34.

40. Vitousek PM, Antonio CM, Loope LL, Westbrooks R. Biological invasions as global environmental change. American Scientist. 1996;84(5):468.

41. Vindenes Y, Langangen O. Individual heterogeneity in life histories and eco-evolutionary dynamics. Ecol Lett. 2015;18(5):417–32. Epub 2015/03/27. doi: 10.1111/ele.12421. PubMed PMID: 25807980.

42. Trauernicht C, Ticktin T. The effects of non-timber forest product cultivation on the plant community structure and composition of a humid tropical forest in southern Mexico. Forest Ecology and Management. 2005;219(2):269–78.

